# Fast unconscious processing of emotional stimuli in early stages of the visual cortex

**DOI:** 10.1101/2021.07.21.453248

**Authors:** Luis Carretié, Uxía Fernández-Folgueiras, Fátima Álvarez, Germán A. Cipriani, Manuel Tapia, Dominique Kessel

**Author notes:** Corresponding author: Luis Carretié, Facultad de Psicología, Universidad Autónoma de Madrid, 28049 Madrid, Spain.

## Abstract

Several cortical and subcortical brain areas have been reported to be sensitive to the emotional content of subliminal stimuli. However, the timing of these activations remains unclear. Our scope was to detect the earliest cortical traces of visual unconscious processing by recording event-related potentials (ERPs) from 43 participants. Subliminal spiders (emotional) and wheels (neutral), sharing similar low-level visual parameters, were presented at two different locations (fixation and periphery). The differential (peak to peak) amplitude from CP1 (77 milliseconds from stimulus onset) to C2 (100 milliseconds), two early visual ERP components originated in V1/V2 according to source localization analyses, was analyzed via Bayesian and traditional analyses. Spiders elicited greater CP1-C2 amplitudes than wheels when presented at fixation. This fast effect of subliminal stimulation -not reported previously to the best of our knowledge-has implications in several debates: i) the amygdala cannot be mediating these effects, ii) latency of other evaluative structures recently proposed, such as the visual thalamus, is compatible with these results, iii) the absence of peripheral stimuli effects points to a relevant role of the parvocellular visual system in unconscious processing.

## Introduction

Our nervous system unconsciously and continuously monitors the environment so salient stimuli -which are, by definition, emotional-may be detected preattentively or, in other words, without consuming controlled, limited brain resources (Carretié, 2014). Such unconscious detection requires the access of the emotional visual input to perceptual brain mechanisms -so it can be identified as a function of its shape, color, motion pattern, etcetera-as well as to evaluative structures -so it is marked as emotional/salient if pertinent-. At this respect, Affective Neuroscience has revealed several brain areas involved in both evaluation and perception to be modulated by the emotional content of subliminal stimulation. Among the former, the activity of the amygdala increases in response to unconsciously perceived emotional stimuli -mainly faces-as compared to neutral (see reviews by Diano et al., 2017; Öhman, 2002). Other brain structures often linked to emotional evaluation, such as the insular or the anterior cingulate cortices, have shown a similar behavior as well (see a meta-analysis in Brooks et al., 2012). Among perceptual structures, the sensitivity of different levels of the visual cortex, from striate (or V1) to various extrastriate areas, such as the fusiform face area, has been also reported (e.g., Axelrod et al., 2015; Brooks, 2012). Despite these data have remarkably increased our knowledge on unconscious processing, several key issues remain unexplored.

Due to the low temporal resolution of functional magnetic resonance imaging (fMRI), the most employed methodology in this field, the timing of unconscious affective processing, and particularly the moment in which the structures mentioned above intervene, is poorly understood. Our main scope is exploring this question and, concretely, detecting the first signs of unconscious visual detection of emotional stimuli. Event-related potentials (ERPs), an electrophysiological manifestation of brain activity with maximal temporal resolution, are especially useful for this purpose, but have been scarcely employed yet. Their main drawback is that, contrary to fMRI, they cannot detect subcortical activity. However, and as explained later, cortical activity may indirectly -but reliably-inform on certain aspects of subcortical activity. Fortunately, most of visual cortex activity is detectable by ERPs so, more specifically, we aim at studying the first visual cortex traces of subliminal emotion detection using this neural recording methodology. Several studies report increased amplitudes in certain ERP visual components around 150-170 ms in response to subliminal facial and non-facial emotional stimuli. As regards facial stimuli, these effects have been reported for the N170 component of the ERPs (e.g., Pegna et al., 2008; Smith, 1012; Zotto & Pegna, 2015; but see Kiss & Eimer, 2008). Face processing areas including the fusiform face area, located in the ventral visual cortex, contribute to the generation of N170 (Deffke et al., 2007; Itier and Taylor, 2004; Sadeh et al., 2010). With respect to non-facial emotional subliminal stimuli results are scarcer. However, greater amplitudes to spiders than to non-negative stimuli have been also observed at 150 ms, the extrastriate visual cortex contributing to generate this activity (Carretié et al., 2005).

Importantly, several ERP components reflecting earlier visual processing take place in short latencies, namely C1, P1p and C2. Whereas their latency varies as a function of the physical characteristics of the stimuli, their spatial location, or the scalp region in which they are recorded, these three components consistently appear before 150ms and two of them, C1 and (often) P1p, before 100ms (Capilla et al., 2016; Di Russo et al., 2012). Their origin also signals these three components as the earliest manifestation of cortical visual processing. Thus, C1 and C2 are mainly originated in the primary visual cortex -or V1- (Capilla et al., 2016; Di Russo et al., 2012; but the contribution of early stages of extrastriate cortex such as V2 and V3 may not be discarded: Ales et al., 2010). On the other hand, P1p is originated in dorsal areas of the visual extrastriate cortex according to these studies. Despite C1 and P1p (C2 has not been explored at this respect) have been reported to be sensitive to the emotional content of supraliminal emotional stimuli (C1: Acunzo et al., 2019; Eldar et al., 2010; Pourtois et al., 2004; P1p: Carretié & Ruiz-Padial, 2016; Holmes et al., 2008; Luo et al., 2010), their sensitivity the emotional content of subliminal stimuli has not been studied yet, to the best of our knowledge. This issue would be relevant at several cerebral levels. At the cortical level, the timing of unconscious processing is less understood than that of conscious processing. Indeed, the first sign of conscious visual processing is well stablished in the literature: the visual awareness negativity (VAN), an ERP component ranging from 150 to 300ms, approximately, and originated in extrastriate ventral areas of the visual cortex (see reviews in Förster et al., 2020; Jiménez et al., 2020). However, first visual cortex traces of unconscious processing remain unexplored.

At the subcortical level, some issues may be also clarified thanks to the temporal resolution of ERPs. One of them is the role of the amygdala in unconscious processing. As indicated, this structure is consistently reported as being sensitive to the emotional content of subliminal stimulation (Brooks et al., 2012; Diano et al., 2017; Öhman, 2002). Some studies have reported the amygdala to respond before visual cortex to subliminal emotional faces (Bayle et al., 2009). In relation to this, the amygdala’s role as an early detector or evaluator of emotional stimuli -not only subliminal-capable of modulating the subsequent activity of the visual cortex has been often postulated (e.g., Framorando et al., 2021; Krolak-Salmon et al., 2004; Sabatinelli et al., 2009; Vuilleumier, 2005). Whether this previous evaluation of the amygdala is preceptive for the visual cortex activity to be modulated by emotional stimulation is, however, unclear. The shortest response of the amygdala to emotional faces reported so far is produced at 74 ms and more than 100ms later in response to emotional non-facial stimuli, according to intracranial recordings (Méndez-Bértolo et al., 2016). Taking into account conduction velocity in middle-range neurons in humans, information from amygdala would reach V1 or V2 at ~95ms and ~200ms in the case of facial and non-facial emotional stimuli, respectively (Carretié et al., 2021). Indeed, the amygdala has been postulated as a key piece of the neural circuitry involved in social behavior and as especially responsive to facial information (e.g., Amaral, 2003; Wang et al., 2014). Therefore, along with time, the facial or non-facial nature of emotional stimuli seems critical in this field. Non-facial emotional stimuli (e.g., harmful animals, food, etc.) may have similar or even more dramatic evolutionary consequences in some circumstances, so they should also be susceptible to being processed unconsciously. However, up to date neuroscience has provided scarce data on the processing of subliminal non-facial emotional stimuli.

The present study explored the earliest ERP visual components presenting non-facial emotional and neutral stimuli using a backward masking task. If a modulation in C1, P1p or C2 is observed, several conclusions may be extracted on the visual cortical (when and which striatal and/or extrastriatal areas are involved, if any) and subcortical roles (any significant effect of emotion in these three components would rule out the preceptive involvement of the amygdala, or other evaluation structures presenting even longer latencies, in unconscious processing. Stimuli consisted of spider and wheel silhouettes. This format was selected since presenting Gestalt characteristics such as closed contours or compact shape (as is the case of silhouettes) are optimal to increase the response of contour-sensitive neurons present in V1 and V2 (e.g., Ko & von der Heydt, 2018). Additionally, spiders are among the top five most feared animals (Gerdes et al., 2009), and they cause the most prevalent phobia related to animals (Jacobi et al., 2004). Subliminal stimuli were presented both at fixation and in the periphery in order to detect possible biases of unconscious processing towards the magnocellular visual processing system (more involved than parvocellular in peripheral processing) activity, as previously proposed (Crick & Koch, 2003). Additionally, this foveal or peripheral presentation helped us to identify C1, C2 and P1p, since they behave differently as a function of the spatial location of stimulation (Capilla et al., 2016).

## Methods

### Participants

46 individuals participated in this experiment, although data from only 43 of them could eventually be analyzed to guarantee an optimal minimum number of trials per condition, as explained later (40 women, age range of 18 to 25 years, mean=19.58, SD=1.33). The study had been approved by the Universidad Autónoma de Madrid’s Ethics Committee. All participants were students of Psychology, provided their informed consent according to the Declaration of Helsinki, and received academic compensation for their participation. They reported normal or corrected-to-normal visual acuity.

### Stimuli and procedure

Participants were placed in an electrically shielded, sound-attenuated room. They were asked to place their chin on a chinrest maintained at a fixed distance (40 cm) from the screen (VIEWpixx®, 120 Hz) throughout the experiment. A backward masking experimental paradigm was employed (Figure 1). Twenty emotional (spiders) and 20 neutral silhouettes (wheels), all in black color over a grey background, were employed as probe stimuli. Spiders are assessed as negatively valenced stimuli by relatively large samples in emotional picture databases (e.g., IAPS: Lang et al., 2005; EmoMadrid: Carretié et al., 2019). In order to test whether spider silhouettes were also efficient as negatively valenced stimuli, and wheels as neutral, these stimuli were previously evaluated by an independent sample of 447 participants (397 women, mean age=19.51, SD=1.46) who rated their emotional valence through a 7-point Likert scale that ranged from “very negative” (1) to “very positive” (7). Spiders were rated as negative (mean=1.704, standard error of means [SEM]=0.038) and wheels as neutral (i.e., in the intermediate values of the scale: mean=3.918, SEM=0.030). Differences between both stimuli were strongly significant (F(1,446)=2557.289, p<0.001, □^2^_p_=0.852).

**Figure 1.**
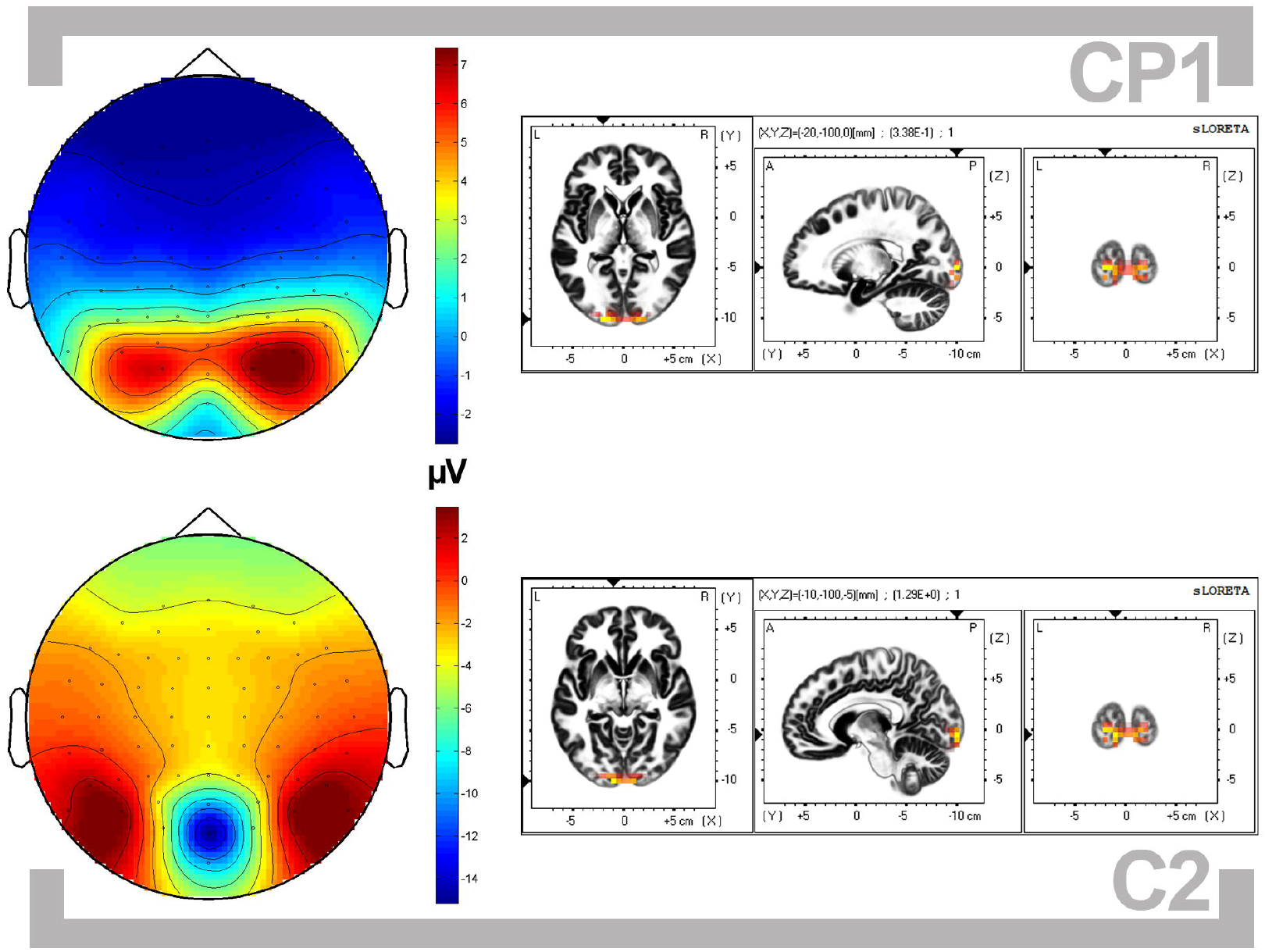
Schematic representation of the stimulus sequence. One of the exemplars of spider and wheel probes are depicted, as well as on mask showing the line in the upper half and another in the lower.

The size of probe stimuli (figure + ground) was 14° × 14° width. Details on the physical characteristics of the spiders and the wheels (i.e., figure surface against background and spatial frequencies), as well as stimuli themselves, are provided in EmoMadrid (www.psicologiauam.es/CEACO/EmoMadrid.htm). Differences between spiders and wheels regarding both spatial frequencies and figure vs ground surface were not significant (see details in the link above). Each spider and wheel appeared 4 times in random order in one of the two locations depicted in Figure 1: at fixation or at the lower visual field. This resulted in 80 trials per emotional category and location, and the total number of trials was 320 (80 × 2 categories × 2 locations). Each probe stimulus, whatever its location, was displayed on the screen for 8.33 ms. We selected this peripheral location since early ERP visual components have previously shown to be more sensitive to emotional stimuli similar to those employed here when they appear in the lower visual field than when presented in other locations of the visual scene (Carretié et al., 2020). Additionally, the lower field is anatomically overrepresented in V1 (Burkhalter et al., 1986), so stimuli in this area tend to elicit greater amplitudes in these components (Capilla et al., 2016).

Immediately after the offset of the probe, a mask appeared for 83.33ms. It consisted of two adjacent squared boxes, each with the same size of probes. The boxes were presented over both possible locations of the probes whatever the actual location of the just presented probe was (Figure 1). The two boxes forming the mask were always the same and consisted of random, colored and unrecognizable composites of small squared fragments of two of the probe pictures. Since both boxes appeared contiguously, they formed a rectangular, perceptually unitary mask. One of the boxes (or mask halves) included a vertical black line (7.7° × 1.15°, centered in that box: Figure 1). The line appeared 50% of trials at the upper box/half and another 50% at the lower in each experimental condition (spider fixation, spider periphery, wheel fixation, wheel periphery). To ensure active, endogenous attention to both locations, participants were asked to press a button if the line appeared in the upper half and a different button if it appeared in the lower. The intertrial interval (i.e., between the mask offset and the next probe onset) was 1500ms. Participants were instructed to look at the fixation dot at the center of the screen all the time, which was marked with a grey circle (0.3° radius) during the interstimulus intervals. The stimulus sequence was divided into two blocks of four minutes approximately with a brief rest period in between.

### Recording and pre-processing

Electroencephalographic (EEG) activity was recorded using an electrode active cap (Biosemi) with Ag-AgCl electrodes. Sixty-four electrodes were placed at the scalp following a homogeneous distribution and the international 10-20 system. The EEG signal was preamplified at the electrode. Following the BioSemi design, the voltage at each active electrode was recorded with respect to a common mode sense (CMS) active electrode and a Driven Right Leg (DRL) passive electrode, replacing the ground electrode. All scalp electrodes were referenced offline to the nosetip. Electrooculographic (EOG) data were recorded supra- and infraorbitally (vertical EOG) as well as from the left versus right orbital rim (horizontal EOG) to detect blinkings and ocular deviations from the fixation point. An online analog low-pass filter was set to 104Hz (5^th^ order, CIC filter). Recordings were continuously digitized at a sampling rate of 512 Hz. An offline digital Butterworth bandpass filter of 0.3 to 30 Hz (4^th^ order, zero phase forward and reverse −twopass-filter) was applied to continuous (pre-epoched) data using the Fieldtrip software (http://fieldtrip.fcdonders.nl; Oostenveld et al., 2011). The continuous recording was divided into 1000 ms epochs for each trial, beginning 200 ms before the probe stimulus onset. The inevitable lag between the marks signaling stimuli onsets (or ‘triggers’) in EEG recordings and its actual onset in the screen was measured employing a photoelectric sensor as described in https://www.youtube.com/watch?v=0BPwcciq8u8,and corrected during pre-processing.

EEG epochs corresponding to trials in which participants responded erroneously or not responded in the task (see the previous section) were eliminated. Blinking-derived artifacts were removed through an independent component analysis (ICA)-based strategy (Jung et al., 2000), as provided in Fieldtrip. After the ICA-based removal process, a second stage of visual inspection of the EEG data was conducted to manually discard trials in which any further artifact, ocular (horizontal or vertical motion) or other type, was present. This automatic and manual rejection procedure led to the average admission of 60.674, 60.279, 60.046, 58.279 (respectively for spider fixation, spider periphery, wheel fixation, wheel periphery; SDs: 6.823, 6.445, 7.101, 6.185). The minimum number of trials accepted for averaging was 50 trials per participant and condition (i.e., each category presented in each location). Data from three participants were eliminated since they did not meet this criterion.

### Data analysis

Behavioral analyses comprised the performance in the line detection task as well as tests of subliminality (both subjective and objective). Regarding performance in the task, errors and reaction times (after removing outliers: ±3 standard deviations) were submitted to repeated-measures ANOVAs introducing Emotion of the probe (spiders, wheels) and its Location (fixation, periphery) as factors. The subjective test of subliminality consisted in asking participants whether they saw “anything apart from the mask and the line”. This question was asked verbally and participants were free to respond whatever they considered. The objective test of subliminality consisted in an additional presentation of the same stimuli at the end of the experimental session. The 20 exemplars of spiders and wheels were randomly presented using the same backward masking setting as in the experimental run (the same exposure times of probes and masks, intertrial interval, and screen locations). In this case, however, the line did not appear in any of both mask boxes, and the task consisted in pressing a key if participants saw a wheel or another key if not (spiders were never mentioned across the experiment to avoid any supraliminal information on the emotional probe and to prevent “contamination” of other potential participants by transmitting this information). The sensitivity index (d’) was computed on the performance of this final task.

As regards ERPs, the first analytic task was identifying and quantifying relevant ERP components (C1, P1p, C2). As later described and discussed, P1p and C1 overlapped in time and manifested as a single “fused component” that shared typical characteristics of both (probably reflecting overlapped processing of probe and mask), so it will be labeled CP1 hereafter. CP1 and C2 were quantified by measuring their peak amplitudes (positive and negative, respectively) within their corresponding windows of interest (72 to 82ms and 95-105ms, respectively). Additionally, and in order to better characterize these components and to localize the processes underlying them within the visual cortices, the Loreta localization algorithm (eLoreta version: Pascual-Marqui, 2007) was computed on these amplitudes.

The experimental effects on these peak amplitudes were analyzed following a two-step procedure. First step was of exploratory nature due to the absence of previous data on the behavior of the earliest ERP visual components in response to subliminal emotional (nonfacial) stimulation. It consisted of Bayesian paired samples T-tests using JASP software (JASP Team, 2020) with the default Cauchy prior (scale 0.707, which represents a medium effect size). Importantly at this respect, Bayesian analyses might not require multiple comparison corrections during statistical tests (Gelman et al., 2012), even when running over 500,000 contrasts (fMRI voxels: Han & Park, 2018). Bayes factors (BF) were computed in relevant scalp channels (i.e., those actually presenting CP1 and C2 components) in order to contrast spider vs wheel amplitudes when presented at fixation, on one hand, and when presented at the periphery, on the other. BFs allowed to assess strength of evidence in favor of the alternative hypothesis (H1: S>W in the case of CP1 or S<W in the case of C2, which had positive and negative polarity, respectively) over the null hypothesis (H0: S=W) in all analyses. A BF_10_ (i.e., informing on the H1>H0 probability) over 3 represents substantial evidence supporting H1 over H0, and less than 1/3, a substantial evidence in the opposite direction: H0 over H1 (Jeffreys, 1939; see also Dienes, 2014).

The second analytical step, of confirmatory nature, consisted of traditional repeated-measures ANOVAs introducing Emotion of the probe (spiders, wheels), its Location (fixation, periphery) and Electrode as factors. Electrodes introduced in this analysis were those detected in the previous Bayesian analyses to present the strongest evidence in favor of H1. Effect sizes in these ANOVAs were computed using the partial eta-square (□^2^_p_) method. Post-hoc comparisons to determine the significance of pairwise contrasts in potential interactions were performed using the Bonferroni correction procedure.

## Results

Individual behavioral and ERP data are openly available at https://osf.io/afq5r/. In the case of ERPs, data are provided in the form of a four-dimension matrix with a size of 43 participants × 4 conditions × 513 data points × 64 EEG recording channels. Table 1 shows means and standard deviations of behavioral and neural parameters in each of the experimental conditions.

**Table 1.** Means and standard error of means (in italics) of error rate, reaction times and CP1-C2 peak to peak amplitude (average of recordings at C6, CP4, P6, P8, PO8 electrodes). See the main text for more details.

|  | Spider fixation | Spider periphery | Wheel fixation | Wheel periphery |
| --- | --- | --- | --- | --- |
| **Error rate (0 to 1)** | 3.95E-04 | 4.17E-04 | 4.84E-04 | 4.75E-04 |
|  | *5.84E-05* | *5.46E-05* | *7.02E-05* | *5.73E-05* |
| **Reaction time (ms)** | 403.610 | 405.113 | 404.732 | 404.721 |
|  | *7.691* | *8.274* | *8.108* | *8.094* |
| **CP1-C2 amplitude (µV)** | 2.539 | 1.668 | 1.935 | 1.755 |
|  | *0.631* | *0.594* | *0.575* | *0.603* |

### Behavioral analyses

As shown in Table 2, ANOVAs on behavioral performance in the line detection task did not yield any significant effect of Emotion of the probe, its Location, or their interaction, neither regarding errors nor reaction times (all p>0.46).

**Table 2.** Main outputs of ANOVAs on errors and reaction times (RTs) in the line detection task for both analyzed factors: Emotion of the probe (neutral, negative), Location of the probe (fixation, periphery) and their interaction. Df=degrees of freedom.

|  | Emotion |  |  |  | Location |  |  |  | Interaction |  |  |  |
| --- | --- | --- | --- | --- | --- | --- | --- | --- | --- | --- | --- | --- |
| | F | df | p | $\eta^2_p$ | F | df | p | $\eta^2_p$ | F | df | p | $\eta^2_p$ |
| **Errors** | 3.169 | 1,42 | 0.082 | 0.07 | 0.021 | 1,42 | 0.884 | 0.001 | 0.268 | 1,42 | 0.607 | 0.006 |
| **RTs** | 0.19 | 1,42 | 0.665 | 0.005 | 0.419 | 1,42 | 0.521 | 0.01 | 0.549 | 1,42 | 0.463 | 0.013 |

The subjective test of subliminality consisted, as indicated, in asking participants -at the end of the experimental session-whether they saw something apart from the mask. Thirty-seven out of the 43 participants responded “no”, and six reported seeing some element unrelated to spiders: “circles” (n=2), “something dentate” (n=1), “maybe a wheel” (n=1), “something black” (n=1), and “a helix” (n=1). This subjective tests was complemented by the objective test, which consisted in the forced-choice task described in Data Analysis and the resulting sensitivity index (d’). Average d’ in the whole sample^1^ was close to zero (8.123*10^−17^, SD=0.5911). As may be appreciated in the supplemental link provided above, except in one case which presented moderate sensitivity (d’=2.300), all d’ were below 1.

### ERPs: identification and characterization of components

Figure 2 shows a selection of grand averages after subtracting the baseline (prestimulus) activity from each ERP. These grand averages correspond to medial and lateral parietooccipital areas, where the relevant visual components were most prominent. As may be appreciated, the two most conspicuous components in early latencies are a positive deflection peaking at 77 ms average (±3) and a negative one at 100ms (±5). Figure 3 shows the topographical distribution of both peak amplitudes as well as their neural origin as revealed by eLoreta. This origin was located in V1/V2 in both components (Table 3). As may be appreciated in these figures, the positive deflection presents characteristics of P1p such as the (slightly, in this case) greater amplitude in response to foveal than to low periphery stimulation (C1 is typically characterized by the opposite pattern), or its bilateral distribution (rather than at midline, as usual in C1; see Capilla et al., 2016; Di Russo et al., 2012 regarding C1 and P1p characteristics). But it also presents attributes of C1 as its latency (slightly shorter than typical P1p) and its origin (V1/V2, whereas P1p is originated in dorsal extrastriate visual areas: Capilla et al., 2016; Di Russo et al., 2012). This overlapping is relatively frequent, but it has probably been enhanced by the rapid probe-mask flip and the concurrence of their respective neural effects. Therefore, and as previously advanced, it is being labeled CP1. As regards the negative deflection, its topographical distribution, its polarity, its latency, and its neural origin (V1/V2) converge to identify it as C2 (Capilla et al., 2016; Di Russo et al., 2012).

**Figure 2.**
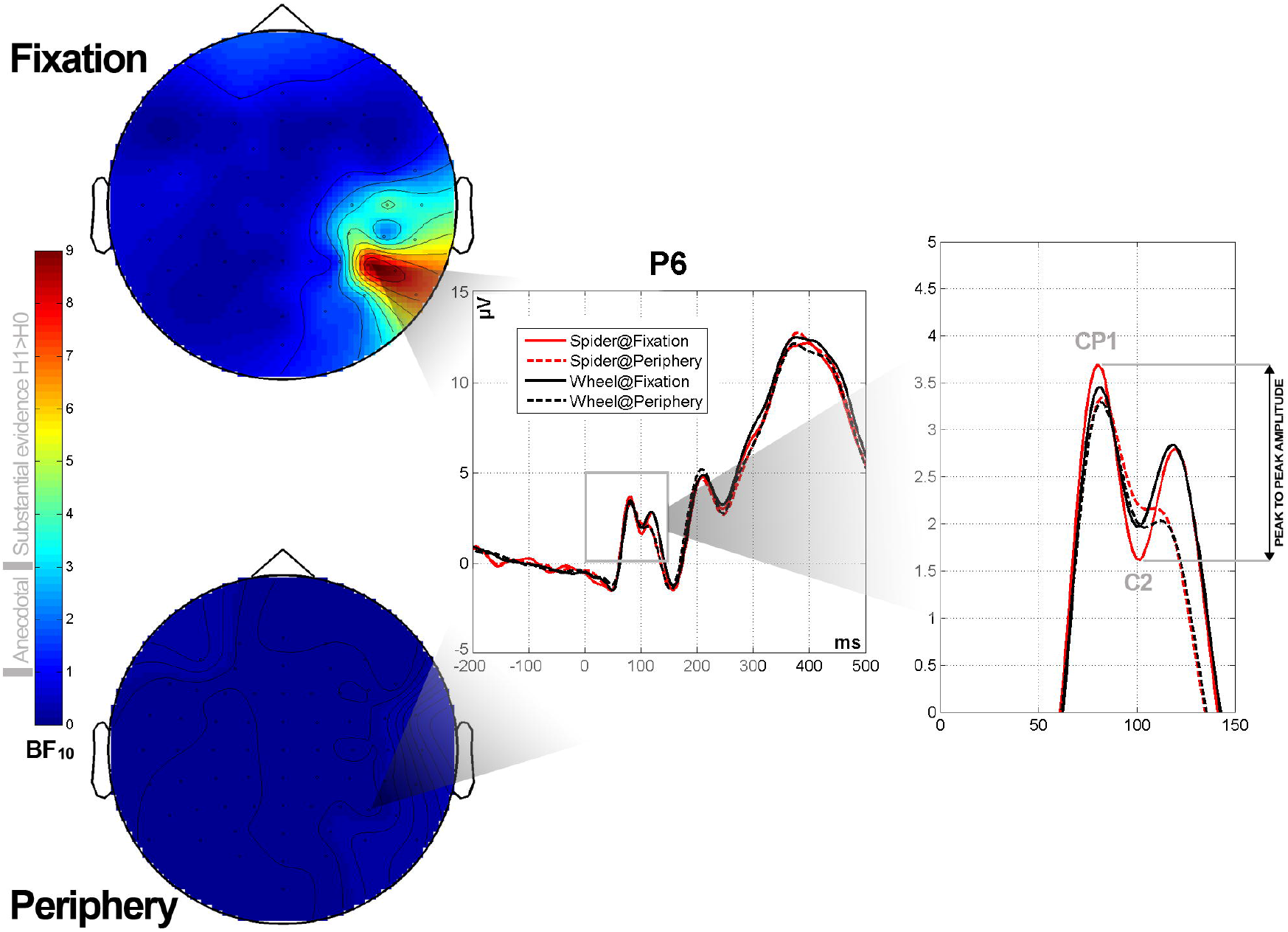
Grand averages (only the −200 to 500 portion of the epoch is depicted) corresponding to five representative sites of the parietal-occipital region, where relevant visual components (CP1 and C2) are prominent.

**Figure 3.**
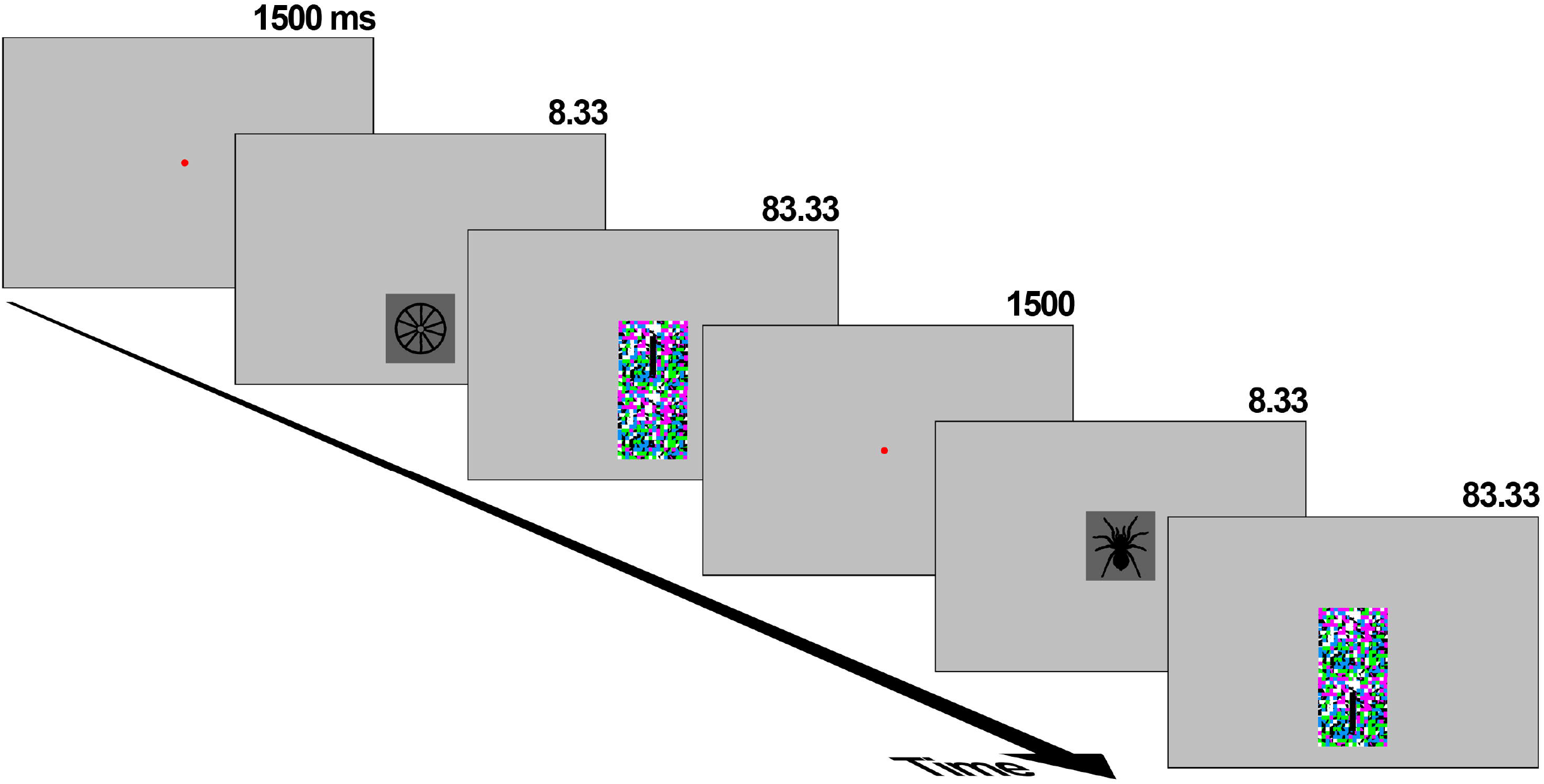
Topographical distribution of CP1 and C2 peak amplitudes (left), and their source as provided by eLoreta (right).

**Table 3.** Main source provided by eLoreta for CP1 and C2 amplitudes.

| ERP component | X,Y,Z (MNI coordinates) | Brodmann area | Subregion |
| --- | --- | --- | --- |
| CP1 | -20 , -100 , 0 | BA18 (0 mm) | V2 |
|  |  | BA17 (7 mm) | V1 |
| C2 | -10 , -100 , -5 | BA17 (0 mm) | V1 |
|  |  | BA18 (5 mm) | V2 |

### ERP: analyses on the experimental effects

A relevant pattern of present ERPs, already visible in the grand averages, is that CP1 to C2 peak-to-peak amplitude appears to be especially sensitive to the experimental manipulation (Figure 4, right). Bayesian, exploratory T-test analyses (see Data Analysis) were carried out on these CP1-C2 peak to peak amplitudes (additional exploratory analyses on CP1 and C2 amplitudes by separate found no evidence to support H1 -Spider>Wheel-since all BF _10_<1.05; see “supplementary information” at https://osf.io/afq5r/). Figure 4 (left) graphically shows the magnitude of Bayesian factors (BF_10_), and Table 4 summarizes the main outputs of all tests. As may be appreciated, strong evidence in favor of H1 (greater CP1-C2 amplitude for spiders than for wheels) is observed in lateral parietal areas, but only when probes were presented at fixation. Concretely, CP1-C2 peak to peak amplitudes in response to probe stimuli presented at fixation yielded BF_10_>3 in electrodes at C6, CP4, P6, P8, PO8 (BF_10_=4.189, 4.105, 8.766, 8.251, 3.178, respectively). Spider vs wheel amplitudes yielded BF_10_<1 in all electrodes (Figure 3 and Table 4).

**Figure 4.**
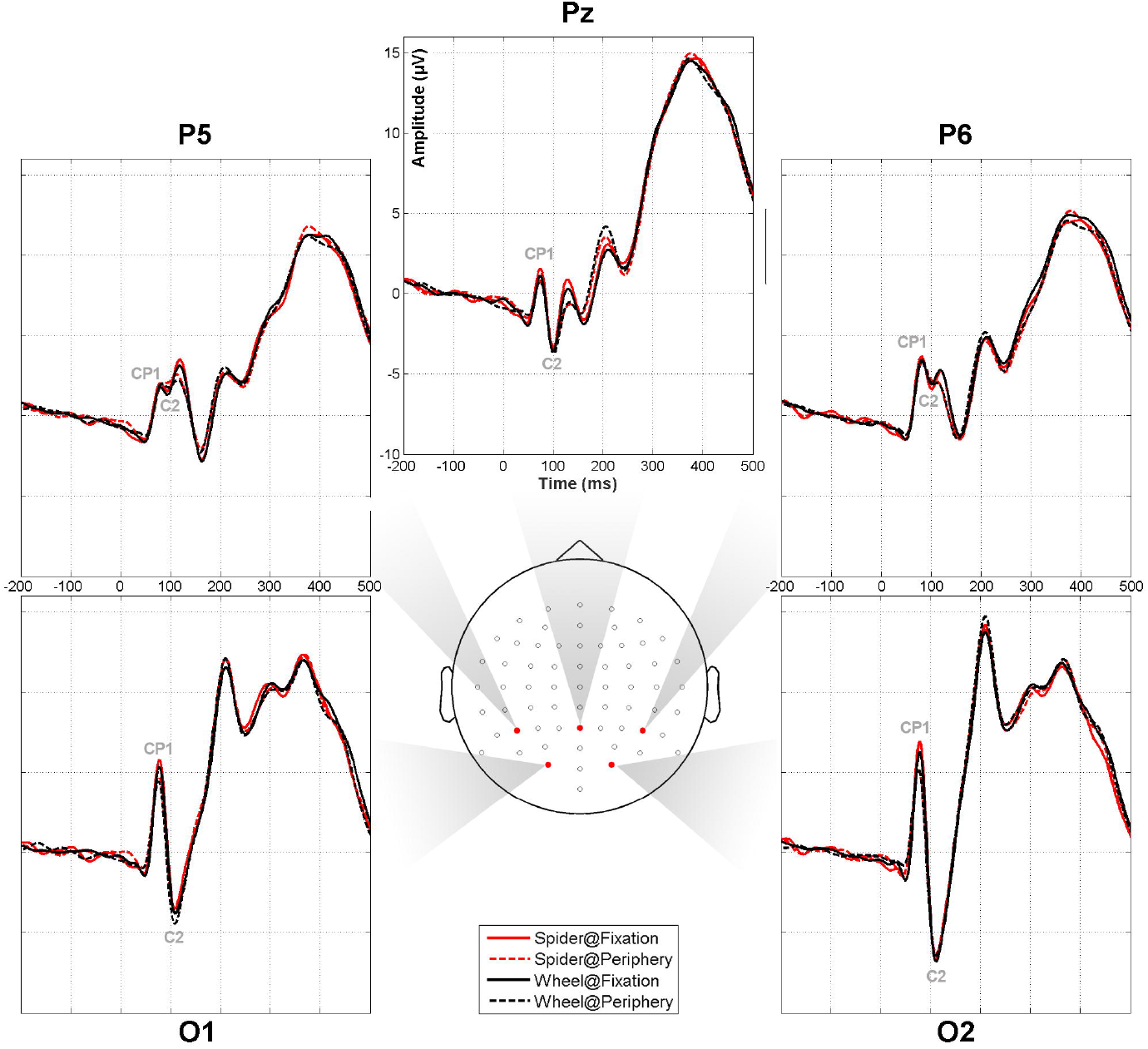
Left: representation, in the form of scalp maps, of Bayesian factors describing the likelihood of data on H1 (spider>wheel) over H0 (spider=wheel) (BF_10_) computed through Bayesian Paired Samples T-Test on the CP1-C2 peak to peak amplitude both when the probe stimuli were presented at fixation and in the periphery. Missing electrodes (see Table 3) were removed and interpolated for this representation. Right: grand averages at one representative electrode of the region showing the greatest BF10 values, with the portion involving CP1 and C2 enlarged to illustrate the peak to peak amplitude quantification procedure.

**Table 4.** Bayesian factors regarding the likelihood of data on H1 (spider>wheel) over H0 (spider=wheel) - BF_10^-^_ computed through Bayesian Paired Samples T-Test using JASP (Jasp Team, 2020) on the CP1-C2 peak to peak amplitude both when the probe stimuli were presented at fixation and in the periphery. Missing electrodes (n=10) are those not presenting any deflection corresponding to CP1 and C2 components (peak to peak amplitude being incomputable).

| Electrode | Fixation | Periphery | Electrode | Fixation | Periphery |
| --- | --- | --- | --- | --- | --- |
| Fp1 | 1.045 | 0.115 | AF8 | 0.319 | 0.089 |
| AF7 | 0.358 | 0.207 | AF4 | 0.187 | 0.11 |
| AF3 | 0.36 | 0.118 | AFz | 0.309 | 0.104 |
| F1 | 0.62 | 0.096 | Fz | 0.258 | 0.107 |
| F3 | 0.405 | 0.078 | F2 | 0.324 | 0.101 |
| F5 | 0.361 | 0.103 | F4 | 0.585 | 0.126 |
| F7 | 0.108 | 0.11 | F6 | 0.316 | 0.121 |
| FC5 | 0.742 | 0.104 | F8 | 0.281 | 0.062 |
| FC3 | 0.45 | 0.082 | FT8 | 1.571 | 0.307 |
| FC1 | 0.349 | 0.082 | FC6 | 1.034 | 0.152 |
| C1 | 0.458 | 0.082 | FC4 | 1.197 | 0.109 |
| C3 | 0.338 | 0.074 | FC2 | 0.472 | 0.122 |
| CP3 | 0.257 | 0.064 | FCz | 0.374 | 0.101 |
| CP1 | 0.423 | 0.077 | Cz | 0.399 | 0.097 |
| P1 | 0.249 | 0.083 | C2 | 1.009 | 0.115 |
| P3 | 0.259 | 0.064 | C4 | 2.644 | 0.135 |
| P5 | 0.283 | 0.06 | C6 | 4.189 | 0.138 |
| PO7 | 0.167 | 0.069 | CP6 | 2.291 | 0.145 |
| PO3 | 0.246 | 0.081 | CP4 | 4.105 | 0.106 |
| O1 | 0.338 | 0.065 | CP2 | 1.345 | 0.107 |
| Iz | 0.809 | 0.064 | P2 | 0.986 | 0.101 |
| Oz | 0.41 | 0.09 | P4 | 1.727 | 0.132 |
| POz | 0.794 | 0.104 | P6 | 8.766 | 0.111 |
| Pz | 0.348 | 0.096 | P8 | 8.251 | 0.155 |
| CPz | 0.511 | 0.1 | PO8 | 3.178 | 0.177 |
| Fpz | 0.83 | 0.068 | PO4 | 1.583 | 0.142 |
| Fp2 | 0.745 | 0.086 | O2 | 1.657 | 0.107 |

Exploratory analyses were followed by confirmatory analyses, as explained in Data Analysis. To this aim, repeated-measures ANOVAs were computed on CP1-C2 peak to peak amplitudes introducing Emotion of the probe (spiders, wheels), its Location (fixation, periphery) and Electrode (C6, CP4, P6, P8, PO8) as factors. Importantly to our scopes, the Emotion × Location interaction resulted significant (F(1,42)=4.878, p=0.033, ⍰^2^_p_=0.104). Post-hoc comparisons showed significant spider > wheel differences when presented at fixation (p=0.006), but not when presented peripherally (p=0.741). These and other less relevant ANOVA results are shown in Table 5. As may be appreciated, interactions with Electrode were all non-significant pointing to a homogeneous behavior of this group of electrodes.

**Table 5.** Main outputs of ANOVA on CP1-C2 peak to peak amplitudes for both analyzed factors: Emotion of the probe (neutral, negative), Location of the probe (fixation, periphery), Electrode (C6, CP4, P6, P8, PO8), and their interactions. Df=degrees of freedom.

| | **F** | **df** | **p** | $\eta^2_p$ |
| --- | --- | --- | --- | --- |
| **Emotion** | 2.08 | 1, 42 | 0.157 | 0.047 |
| **Location** | 4.862 | 1, 42 | 0.033 | 0.104 |
| **Electrode** | 18.675 | 4, 168 | <0.001 | 0.308 |
| **Emotion x Location** | 4.878 | 1, 42 | 0.033 | 0.104 |
| **Emotion x Electrode** | 0.585 | 4, 168 | 0.543 | 0.014 |
| **Location x Electrode** | 0.788 | 4, 168 | 0.451 | 0.018 |
| **Emo x Locat x Electr** | 0.406 | 4, 168 | 0.813 | 0.006 |

## Discussion

The present study explored the earliest traces of unconscious processing of emotion in the visual cortex by recording ERPs, a temporally agile neural signal. The relatively large sample size ensured an adequate statistical power. Two different analytical strategies (Bayesian and traditional) revealed a short-latency sensitivity (72-105ms) of early stages of the visual cortex (V1/V2) to the emotional content of non-facial stimuli presented subliminally. These results are especially valuable since they were obtained using a backward masking paradigm, in which EEG and MEG have more difficulty to manifest unconscious processing effects (as compared to other paradigms such as binocular rivalry; Axelrod et al., 2015). Additionally, they were obtained using a relatively short probe duration, since spiders and wheels were exposed only during 8ms. Several characteristics of the experimental design may have helped to obtain these early effects for the first time -to the best of our knowledge-. First, the use of spiders, an emotionally potent stimulus, as explained in the Stimuli and Procedure section, may intensify emotion detection mechanisms as compared to other stimuli (including faces). Second, the use of black silhouettes over a lighter background increases the response of contour-sensitive neurons present in V1 and V2, contributing to augment the signal-to-noise ratio of the activity detected at the scalp level. And third, the relatively large number of trials (~60 average per condition, 50 minimum) also contributes to increase the signal-to-noise ratio.

These results have important implications not only at the cortical level, but also at the subcortical. Regarding the former, they reveal certain parallelism with what occurs in response to supraliminal emotional stimulation, since early visual ERP components also show sensitivity to emotional stimuli (C1: Acunzo et al., 2019; Eldar et al., 2010; Pourtois et al., 2004; P1p: Carretié & Ruiz-Padial, 2016; Holmes et al., 2008; Luo et al., 2010). However, the effects are more difficult to detect in this case and, along with the implementation of the issues mentioned in the previous paragraph, have required the quantification of differential CP1-C2 amplitudes (a classical way of computing amplitudes that seems useful in this field) rather than individual component amplitudes. Also at the cortical level, these results contribute to balance the knowledge on the temporal characteristics of unconscious vs conscious visual processing. On the conscious side of the coin, there is a relatively broad consensus that the earliest traces of conscious processing are reflected in VAN, as indicated in the Introduction (Förster et al., 2020; Jiménez et al., 2020). Whereas from this it would follow that earlier visual activity remains in the unconscious domain -although consequent processes may become conscious-, data from studies directly exploring the effects of subliminal information are necessary for this temporal characterization.

Subcortical implications of these results are also relevant, mostly in the emotional field. As also indicated in the Introduction, any early effect of subliminal emotional stimuli, especially if they are non-facial, automatically rules out the involvement of the amygdala in this phase (its involvement in later phases of emotional processing -conscious and unconscious-is clear at the light of existing data). Even if stimuli were faces, the latency of the observed effects (72-105ms) would also be incompatible with the involvement of the amygdala in visual cortex modulation. Indeed, the earliest response of amygdala reported so far is produced at 74 ms, and more than 100 ms later in the case of non-facial emotional stimuli (Méndez-Bértolo et al., 2016). This amygdalar activity would require ~20-25 additional milliseconds to reach the visual cortex, as explained in the Introduction. The probability that the amygdala may be found in future studies to respond significantly earlier than this to visual stimuli is remote due to anatomical and functional reasons detailed elsewhere (Carretié et al., 2021). Relatedly, the probability that other structures postulated as evaluative, such as ventral prefrontal, insular or anterior cingulate cortices intervene at this latency in visual cortex modulation is even more remote, due to their even longer latencies in response to emotional stimuli (Adolphs et al., 2006; Willenbockel et al., 2012). Indeed, present results point to alternative hypotheses regarding brain areas responsible for initial stimulus evaluation, particularly to first order structures (i.e., retinal ganglion cells directly reach them), due to their short processing latency. This is the case of the visual thalamus and other associated subcortical visual nuclei, recently revealed as active processors rather than passive relays in the visual ascending route towards visual cortices, as traditionally considered (Carretié et al., 2021).

Two additional results are worth discussing. On one hand, only spiders presented at fixation elicited enhanced amplitudes as compared to neutral stimuli. Retinal projections to parvocellular and magnocellular layers of the lateral geniculate nucleus decline to a greater extent in the former case with eccentricity (Brown et al., 2005), resulting in a magnocellular bias for peripheral vision. This is a relevant issue since some traditional postulates defend a preferential involvement of the parvocellular system in conscious visual processing and the magnocellular in unconscious processing (Crick & Koch, 2003; Milner & Goodale, 1995). Present results point against this sort of segregation in line with more recent views (Breitmeyer, 2014). On the other hand, although both CP1 and C2 were originated in the cuneus (V1/V2), the CP1-C2 peak to peak effects were observed more laterally and extending towards the parietal cortex. The fact that the experimental effects are not reflected where amplitudes are more prominent is relatively frequent, however. This phenomenon has been scarcely studied, but probably reflects the involvement of additional, secondary sources being responsible for the observed effects. This is even more probable when, as in our study, multiple overlapping are produced: CP1 presents mixed characteristics of C1 and P1, and the peak to peak amplitude further involves the processes underlying C2. Although source reconstruction was not possible for the experimental effects, since not all electrodes were analyzed (those not actually showing CP1 and C2 were discarded), we hypothesize -at the light of the scalp distribution of effects: Figure 4) that dorsal extrastriate areas are probably involved. Some dorsal extrastriate areas, such as MT, receive rapid and direct inputs from the visual thalamus which, as mentioned, has been recently proposed as an early evaluation structure (e.g., from the inferior pulvinar, from lateral geniculate nucleus and from the visual portion of the thalamic reticular nucleus: Carretié et al., 2021).

In sum, present results show the short-latency sensibility of early stages of the visual cortex to the emotional content of unconsciously perceived stimulation. This finding confirms the particular (and adaptive) relevance of emotional stimuli for our nervous system. Several implications derived from this result have been discussed throughout this section. They go beyond the visual cortex level and also beyond emotional processing and would require further research to be confirmed. At this respect, ERPs demonstrate to be a valuable tool in this scarcely explored field, although several manipulations to increase the signal-to-noise ratio of recordings, such as those mentioned previously, seem necessary.

## Acknowledgements

This research was supported by the Ministerio de Ciencia e Innovación (MICINN) (Grant no. PGC2018-093570-B-I00) and the Comunidad de Madrid (Grants no. HUM19-HUM5705 and, in collaboration with the Universidad Autónoma de Madrid, SI1-PJI-2019-00011 and 2017-T2/SOC-5569).

## Footnotes

1 Two out of the 43 participants responded an insufficient number of trials (1 and 6 out of 40) due to their misunderstanding of the forced-choice task instructions (they avoided pressing any button whenever they ignored the response), so their d’ was not computed. In any case, their blank response rates objectively confirm, in an alternative way, the subliminal condition of probes also for these two participants. The rest of participants responded in more than 75% of trials.

## Notes

https://osf.io/afq5r/

### Competing Interest Statement

The authors have declared no competing interest.

